# Phages - bacteria interactions network of the healthy human gut

**DOI:** 10.1101/2020.05.13.093716

**Authors:** Martial Marbouty, Agnès Thierry, Romain Koszul

**Author notes:** corresponding authors, contacts or.

## Abstract

With an estimated 10_31_ particles on earth, bacteriophages are the most abundant genomic entities across all habitats and important drivers of microbial communities. Growing evidence suggest that they play roles in intestinal human microbiota homeostasis, and recent metagenomics studies on the viral fraction of this ecosystem have provided crucial information about their diversity and specificity. However, the bacterial hosts of this viral fraction, a necessary information to characterize further the balance of these ecosystems, remain poorly characterized. Here we unveil, using an enhanced metagenomic Hi-C approach, a large network of 6,651 host-phage relationships in the healthy human gut allowing to study *in situ* phage-host ratio. We notably found that half of these contigs appear to be sleeping prophages whereas ¼ exhibit a higher coverage than their associated MAG representing potentially active phages impacting the ecosystem. We also detect different candidate members of the crAss-like phage family as well as their bacterial hosts showing that these elusive phages infect different genus of Bacteroidetes. This work opens the door to single sample analysis and concomitant study of phages and bacteria in complex communities.

## Introduction

The gut hosts a microbial ecosystem composed of bacteria, archaea, eukaryotic microorganisms and viruses. High-throughput sequencing now allows in-depth investigation of the human gut microbiota^1,2^, unveiling links between the balance of this complex ecosystem with a wide range of human diseases^3^, including autoimmunity^4^ or neurological disorders^5^.

More recently the viral part of the human gut microbiome started to gain greater attention as roles for phages in regulating health emerged either by directly influencing the immune system or by reshaping the bacterial component of the gut^6–9^. Human gut virome is dominated by temperate bacteriophages belonging to the Caudovirales family^10,11^. However, how the microbiome and the virome influence each other’s remains largely unknown, with few analysis characterizing concomitantly host-phage relationships in the gut^12,13^, a key, primary step to understand the impact of phages on microbial population^14^. Phages – bacteria ratio in human gut is another important question and is mainly addressed through quantification of virus-like particles corresponding to free phages produced during lytic cycle^15^. The studies that tackle this question yield values that appear below global bacterial concentrations (between 1 and 0.001)^11,16,17^, suggesting that the intestinal tract is quite different from other ecosystems such as oceans^18^ where phages largely outnumber bacteria. Therefore, new approaches and studies are needed to characterize 1) which phage infects which bacterial species and 2) the relative phage and bacterial concentrations for each phage-host pair.

In recent years, several methods have been developed that are able to identify whether two, or more, DNA molecules are colocalized in the same genome, and conserve this information throughout sequencing steps. These techniques, which include proximity ligation^19–22^ (e.g. meta3C), viral tagging^12,23^ (VT) or single amplified genomes^24,25^ (SAGs) have allowed to capture directly for the first time the global host-phage network in natural environment. We showed that meta3C, which quantified collisions between DNA molecules, allow the recovery of dozens of phages and bacteria genomes and their relationships, from mammalian gut^19,26^.

Here we applied the meta3C proximity ligation technique to reconstruct the whole landscape of phages and their interacting bacteria of 10 healthy human guts microbiota by capturing the DNA – DNA collisions during phages replication inside their bacterial hosts in each sample. Our work results in 6,651 unique host-phage contigs pairing and constitutes the largest host-phage network of the human gut microbiome to date. While half of the detected phages appeared to be dormant lysogenic phages, we characterize a significant proportion of the phage’s population exhibiting a coverage above the one of their corresponding host. We also characterized different crAss-like phages candidates as well as their bacterial hosts, all belonging to Bacteroides. This study can have practical consequences, notably for faecal microbiota transplantation (FMT)^27^ or phage therapy^28,29^.

## Results

### High-quality proximity ligation meta3C libraries of human gut microbiome

Frozen stool samples from 10 healthy adults from the Institut Pasteur biobanque (agreement N18) (Fig. 1a) were processed with meta3C using either HpaII or MluCI as a restriction enzyme^19^. Each meta3C library was sequenced (between 43 and 206 million pair-end reads) and assembled using MegaHIT^30^, resulting in 10 human gut assemblies ranging from 242 Mb to 617 Mb (total size = 3.45 Gb for 1,485,156 contigs) (Methods) (Supplementary Table 1). To assess the amount of long-range 3D DNA contacts in the different meta3C libraries, each read of a pair was aligned against its corresponding assembly. The number of pairs for which both reads aligned unambiguously on two different contigs (mapping quality ≥ 20) was used to compute a “3D ratio”, by dividing it with the total number of pairs of reads that aligned unambiguously within individual contigs (Supplementary Fig. 1 and Supplementary Table 2). 3D ratios using the standard meta3C protocol ranged from 1.92 % to 14.58 %, a variability that results most likely from samples heterogeneities and the restriction enzyme. These ratio are also in the range, if not higher, of recently published dataset that use meta3C related approaches (0,36 and 2.38% from^31,32^ respectively). The relatively limited outcome of these protocols, which include a biotin enrichment step, suggest room for improvement for both public and commercial solutions.

**Figure 1:**
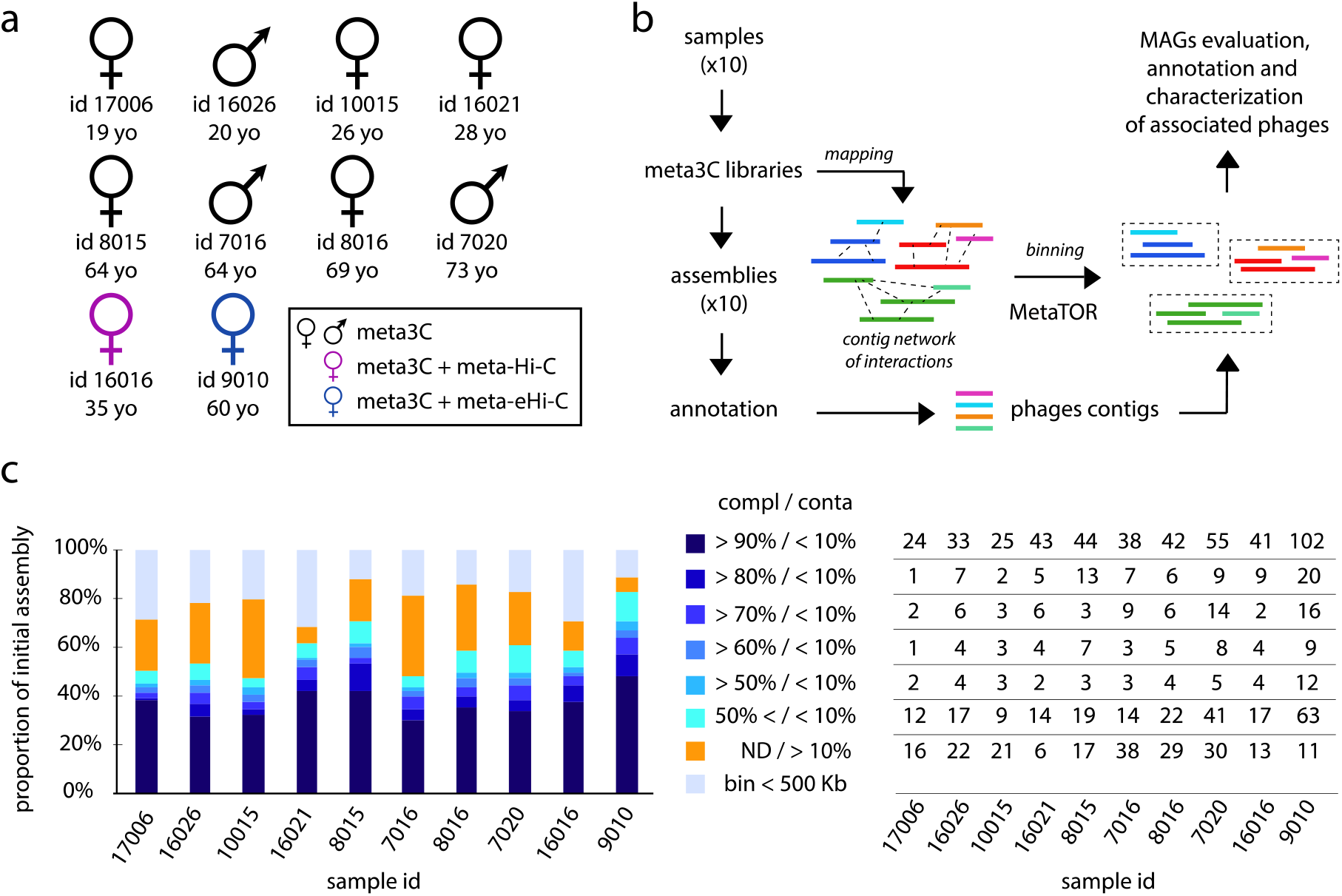
Binning of human gut microbiota using 3C, Hi-C and eHi-C. **a.** Sex, ID and age of the 10 individuals investigated in this study. The colors indicate the proximity ligation protocol used to generate the libraries. **b.** Computational pipeline of PE sequences. **c.** (Left panel) For each individual, proportion the corresponding assembly of the different bins obtained for the 10 fecal samples. The colored proportions represent groups of bins of similar quality (completion and contamination), according to the color code on the right. ND = not determined. (Right panel) For each individual, number of MAGs corresponding to the histogram on the left.

To further the libraries in long-range contacts, we developed two derivatives of meta3C, including one, meta-eHi-C, inspired from other work on bacteria and applied them either on samples #16016 and #9010 (Fig. 1a). In our practice, adding a simple biotin enrichment step to meta3C (meta-Hi-C) only marginally improved the ratio (14.91 % and 5.95 % compared to 4.48 % and 5.15 % for the HpaII and MluCI libraries, respectively). However, the ratios obtained for #9010 using meta-eHi-C were strongly improved for both enzymes (41.22 % and 36.02 % compared to 9.05 % and 7.6 % for the Hpall and MluCI libraries, respectively; Supplementary Fig. 1), suggesting that several times more informative trans contacts between pairs of contigs can be obtained at a similar cost using the meta-eHi-C derivative.

### Binning of human gut microbiome using 3C, Hi-C or eHi-C

We applied metaTOR 26, an iterative network segmentation pipeline, to identify communities of contigs (bins) enriched in trans contacts and corresponding to individual genomes (Fig. 1b) (Methods) (https://github.com/koszullab/metaTOR). For each metagenome the qualities of the metaTOR bins were assessed using CheckM, which quantifies the contamination and completion of pools of contigs with respect to a standard bacterial genome 33. Among the 1,485,156 contigs (3.45 Gb) present in the 10 assemblies, 300,677 (525 Mb) were assigned to a bin ranging in size between 10 and 500 kb (n = 19,122), and 852,876 (2.57 Gb) were found in 1,086 bins containing at least 500 kb and defined as Metagenome Assembled Genome (MAG). MAGs represented between 72 and 91% of all mapped reads depending on samples. CheckM turned up 285 MAGs as “high-quality”, i.e. displaying a completion rate ≥ 90% and contamination rate ≤ 5%, and corresponding most likely to complete bacteria genomes. In addition, 405 MAGs were labelled as “medium quality” (completion ≥ 50% and contamination ≤ 10%) (Fig. 1c) (Supplementary Dataset 1).

We analyzed the contribution of the improved meta-eHi-C protocol to the generation of high-quality MAGs. Indeed, a high 3D ratio does not necessarily mean an enrichment in informative bridges between contigs, as high levels of noise (i.e. ligation events between random restriction fragments) could also increase the ratio. However, all metrics (number of high-quality MAGs, completion, etc.) recovered when comparing the binned meta-Hi-C and meta-eHi-C metagenomic maps with their meta3C equivalent point at a strong enrichment in truly informative contacts (Supplementary Fig. 3; Methods). Bins generated from meta-eHi-C maps showed an improvement in signal-to-noise ratio, with less contacts between them and more contacts within. This is further confirmed by the measure of the modularity^34^ for the two corresponding networks (0.93 for meta-eHi-C compared to 0.86 with meta3C), pointing at a better-defined community structure using the meta-eHi-C network. A comparison between the MAGs retrieved using the two datasets showed that both approaches are highly concordant. The meta3C method results in a slightly noisier signal, which can be overcome by a higher sequencing depth (Supplementary Fig. 2). Therefore, the meta meta-eHi-C protocol is several times cheaper than conventional meta3C or commercial solutions, and should be chosen in future studies.

### Diversity of the human gut microbiota assessed by meta3C

We next generated a phylogenetic tree of all the MAGs whose completion was above 50% and contamination under 20% (Fig. 2) (Methods). A tree was also generated for each individual sample (Supplementary Fig 3). As expected, all trees displayed the taxonomic groups found in a healthy human intestinal tract^35^. To further validate the approach, we show that the abundancy of the characterized MAGs displays a strong correlation with the outcome of taxonomic analysis of the raw reads by the classifier Kaiju^36^ (Supplementary Fig. 4). We then compared the MAGs generated by the meta3C approach to genome references of the GTDB-Tk databases^37^. 394 (~1/3rd) of the 1,086 characterized MAGs and 55 (~1/6th) of the 285 high quality MAGs didn’t show any close similarity to the GTDB-Tk references at a threshold identity of 95% (Average Nucleotide Identity – ANI^38^). These new genomes are proportionally distributed among the different clades, with a majority belonging to Clostridia. Some of these genomes belong to species with an abundancy below ~0.1 % (Fig. 2, Supplementary Fig. 5 and Supplementary Dataset 1), paving the way to future in-depth individual sample analysis. The enhanced meta-eHi-C protocol was able, from 200 million PE reads on a single sample, to identify 81 high-quality genomes including from 20 unknown species. This result demonstrates the robust added value of the approach to any study aiming at investigating complex microbial communities, especially from single samples.

**Figure 2:**
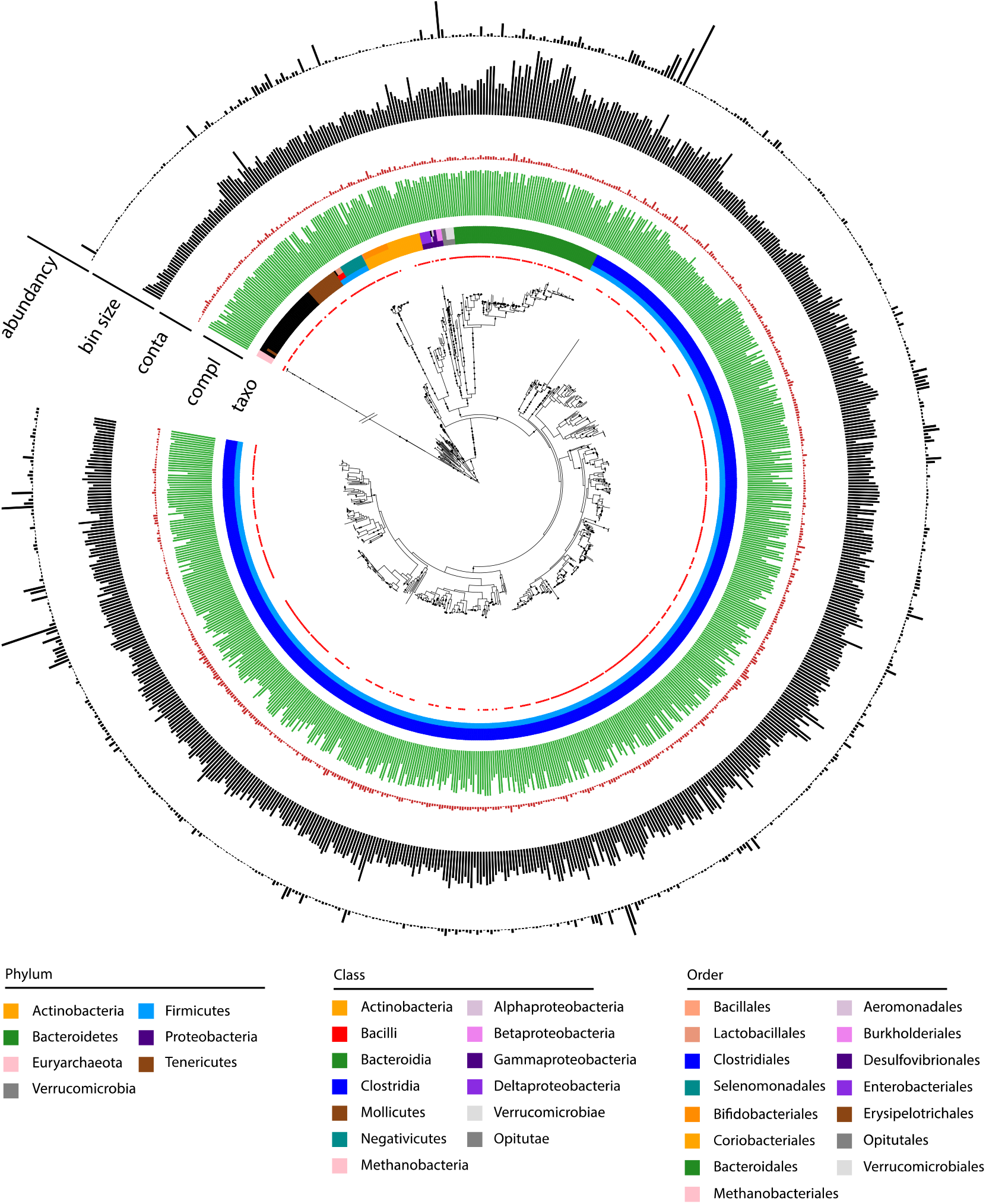
MAGs recovered using Meta3C applied on 10 healthy human gut samples. Phylogenetic tree comprising the 757 reconstructed MAGs with a completion above 50% and a contamination below 20% (751 different species at a threshold of 95% identity). The branch of the Euryarcheota has been cut (//) to resize the tree. Red dots indicate MAGs with a reference in the GTDB-tk database (threshold of homology = 95%). Colors of the three first layers indicate the taxonomy of the MAG attributed by CheckM (from center to periphery: Phylum, Class, Order). Green and red bars in the following circos indicate completion and contamination, respectively. Completion (green) scale: max = 100%; min = 50%. Contamination (red) scale: max = 20 %; min = 0%. Grey bars in the following circos indicate MAG (bin) size (scale: max = 7.07 Mb, min = 766 kb). Grey bars in the last layer indicate MAGS abundancy in the different samples (max = 4.73 %; min = 0.0039 %).

### Phages-hosts network in the human intestinal tract

To assign phages to the different characterized MAGs, we searched for contigs annotated as phages using the VIRSorter^39^ and VIBRANT^40^ programs. Among the 1,485,156 contigs, VIRSorter and VIBRANT identified 4,649 and 7,918 putative phages, respectively (excluding identified prophages), resulting in 9,636 (unique) annotated phages contigs, most of them between 1 to 10 kb. Taxonomic annotations of those phages contigs using DemoVir (https://github.com/feargalr/Demovir) confirmed the high prevalence of phages belonging to the Caudovirales order (families Siphoviridae 59 %, Myoviridae 15%, Podoviridae 2%, Unassigned 22 %). The trans contacts between these phage annotated contigs and the 1,086 MAGs reflect the infection pattern of the bacterial species presents in the 10 samples 19,20. Three classes (A, B, C) of phages were defined (Methods; Fig. 3a; supplementary Fig. 6). First, 6,651 (class A) phages contigs that were unambiguously assigned, based on their 3D contacts, to a single MAG (Supplementary Dataset 4). In total, 931 MAGs out of 1,086 made specific 3D contacts with one, or more, of these contigs. This result suggests that the majority (~69%) of the phages present within the human gut are specific, infecting only one bacterial species. The remaining 2,985 phage annotated contigs may correspond (class B) to phages able to infect several species (n = 2,460), or (class C) to free phages with no contacts to MAGs, or associated to uncharacterized/poorly abundant MAGs (n = 525). Categories B and C phages were excluded from subsequent analysis and saved for future studies (Supplementary Dataset 4). A MAG was associated on average with 7.1 phages contigs (SD = 7.5), but this rate presented high fluctuation: up to 59 contigs were assigned to a single MAG.

**Figure 3:**
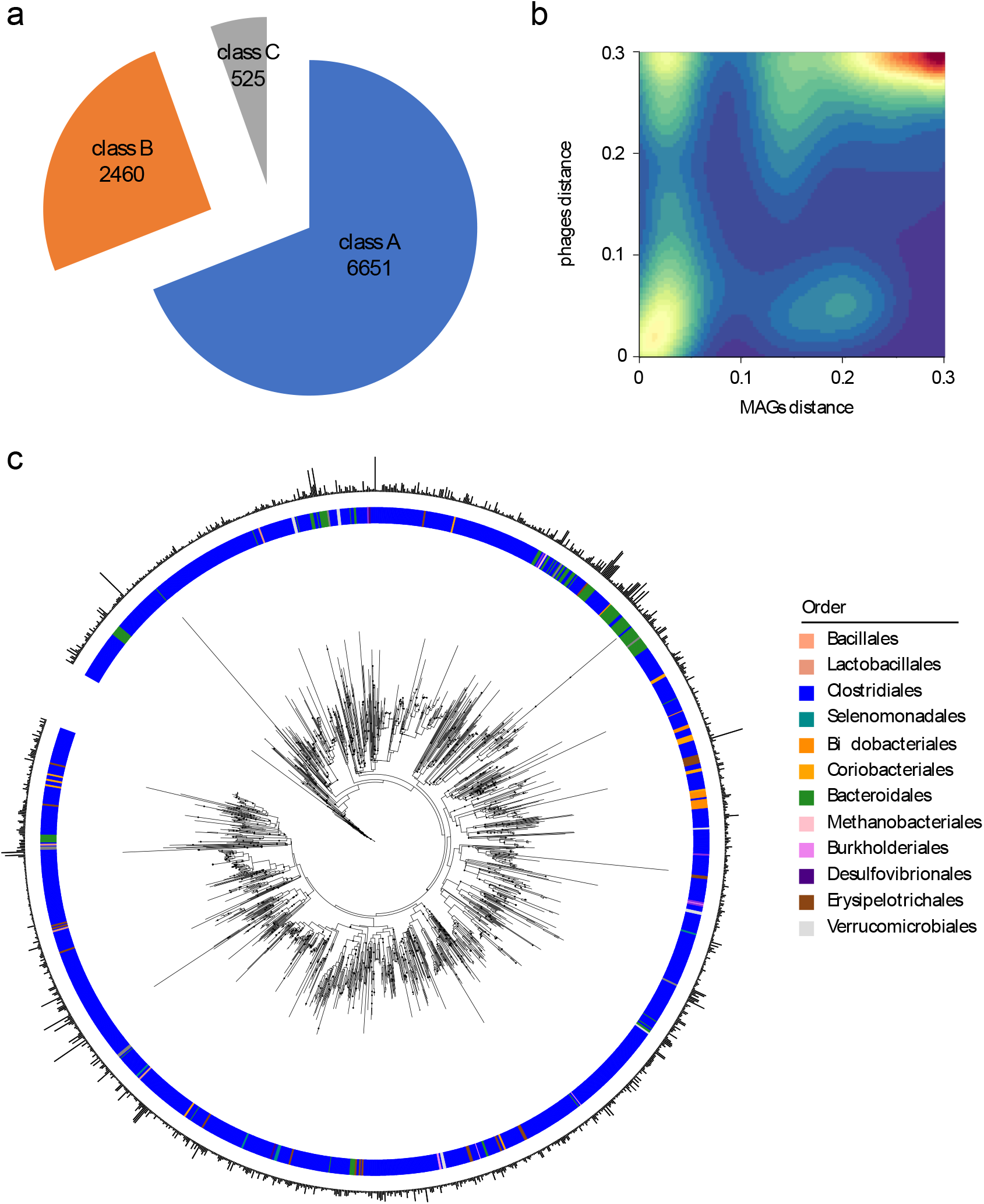
Phages - bacteria network of interactions in human gut. **a.** Pie chart of phages contigs distribution among the different classes (see Methods and Supplementary Fig. 6). **b.** 2D density plot of distances between MAGs and their associated phages. **c.** Phylogenetic tree of phage contigs (n= 2,714). Colored strips in the inner circle indicate the taxonomy of the associated MAG (at the order level). Bar plots in the outer circle indicate the size of the phage contigs (minimum = 5,002 bp; maximum = 211,086 bp).

Of 454 pairs of phages from class A identified as belonging to the same genus (*i.e*. defined as contigs sharing at least 70% identity on 70% of their length^41^), 394 were assigned to a pair of MAGs with the same taxonomic assignment (Supplementary Fig. 7). This result highlights the accuracy of our methods and the value of the phage-host network provided in this study. To further study the infection network generated by our approach, we computed the distance homology between MAGs as well as between all phages contigs using the program Mash which estimates the mutation rate between two sequences directly from their MinHash sketches^42^ (Methods). We then plotted the distance between phages as a function of the distance between their related MAGs and show that homolog phages were associated to homolog MAGs (Fig. 3b, bottom left corner). In addition, while a certain proportion of distantly related phages could occasionally be found infecting homologous MAGs, there was very few instances were related phages would infect non-homologous MAGs (Fig. 3b). This result support the idea that phages are “locally adapted” to their bacterial hosts and do not easily shift from one bacterial species to another^43,44^. Our phages – bacteria network was consistent with CRISPR analysis even if, as expected, only 15 % of the pairing could be cross validated using this computational approach (Methods).

To provide an overview of the phages contigs – bacteria network of the human gut, we computed a phylogenetic tree of the phages (1474 phage contigs included over the 6651) as previously described 45 (Methods). Using both the MAGs taxonomic annotation from CheckM and phage contigs phylogenetic trees, we bridged, using meta3C contacts, the phages and bacteria taxon at the genus level (Fig. 3c) (Supplementary Dataset 4). This dense network, which aims at being refined with future studies, unveiled clusters of phylogenetically distant phage contigs infecting various genus of bacteria. It also shows the existence of clusters of closely related contigs infecting the same genera suggesting, again, specificity of phages from human gut.

### Phages host ratio in the human intestinal tract

The availability of a phages-bacteria network provides new ways to gain insights on the metabolisms of these phages. Indeed, although we ignore the proportion of infected cells in the host population, comparing the read coverages of the phage (contig) and host (MAG) genome can reveal useful information about the former’s behavior (Fig. 4). For the 6,651 phages sequences assigned to a unique MAG, the mean read coverage ratio is 1.26 (SD = 1.78), with a maximum of 56 (Fig. 4a and Supplementary Fig. 8). This distribution is similar to the one of all the contigs contained within our different MAGs. There were nonetheless significant variations among the 10 human individual microbiota, as the average ratio varies from 1 (sample #10015) to 2 (sample #17006) (Supplementary Fig. 8). These variations could reflect differences between individual gut microbiome or through time, and more analysis, larger cohort and time-series sampling are needed to address this question.

**Figure 4:**
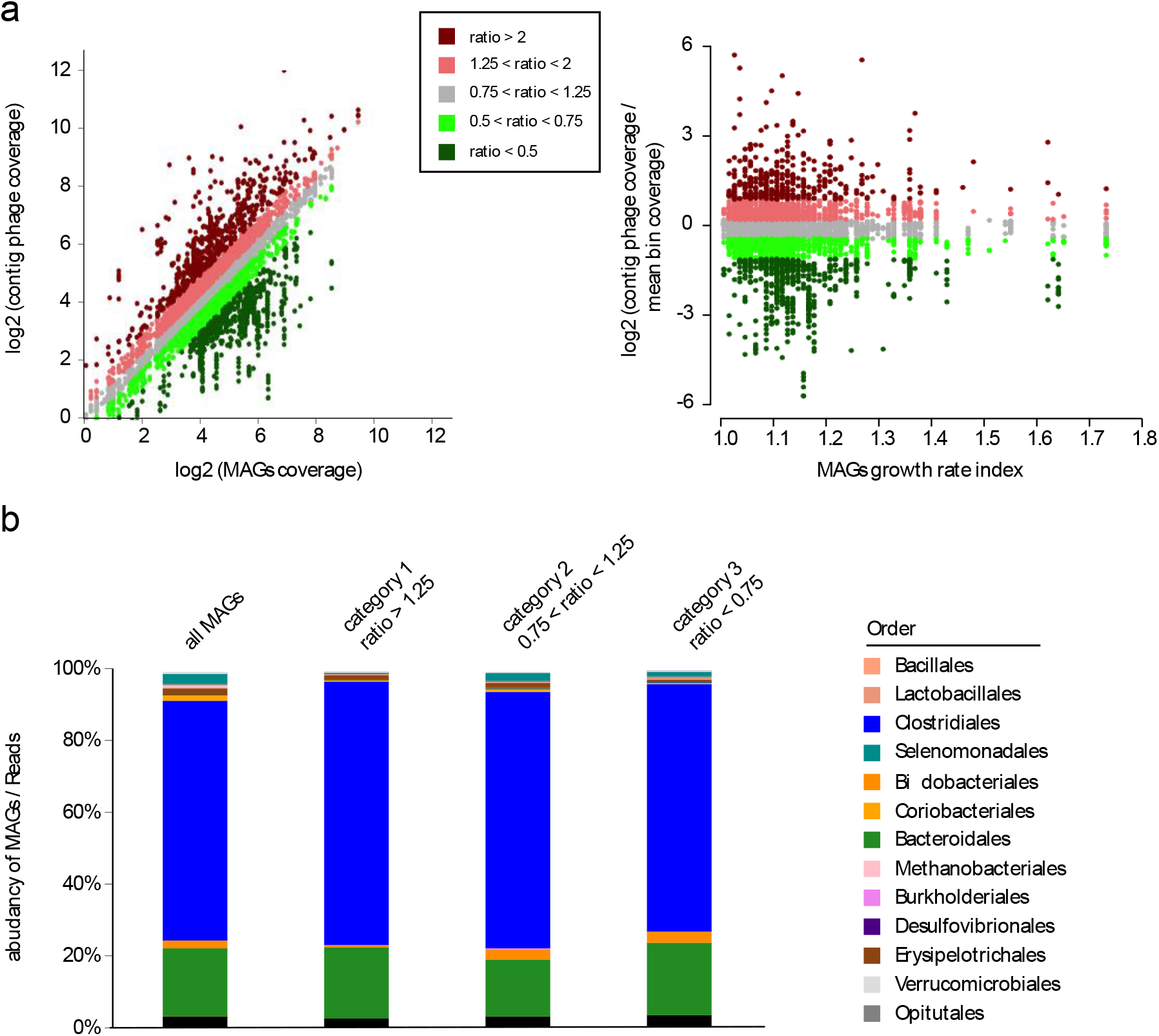
Phages - host ratio in human intestinal tract. **a.** Left panel: plot representing the read coverage of Class A phage contigs as a function of the average read coverage of their associated MAG. Right panel: plot of the ratio of phage contigs coverage vs. MAG mean coverage as a function of the Growth Rate Index calculated using GRID. Dots are colored according to the ratio, with the color code indicated in the box between the graphs. **b.** Bar plot of MAGs taxonomic abundancy for the different categories of their associated phages (category 1 – ratio > 1.25; category 2 – 1.25 > ratio > 0.75; category 3 – ratio < 0.75).

We characterized 3 different categories of phages (Fig. 4a). First, phages displaying a ~1:1 ratio (+/- 25 %) with a MAG genome. This population represents nearly half of the 6,651 phages-host pairs (n = 3,152; 1,444 within +/- 10 %) and could correspond to sleeping prophages (able to reactivate or not) or active phages that, by chance, exhibit the same coverage of their host. Then, we detect 1,650 pairs for which the MAG is significantly more abundant than the associated phage. This profile is consistent with an abortive phage infection cycle, a pseudo – lysogenic cycle or even with prophages not present in the whole host population. Finally, we detect 1,849 pairs corresponding to a higher phage coverage compared to its corresponding MAG (604 exhibit a ratio ≥ 2). Those could represent phages actively replicating and potentially impacting the ecosystem through lytic activity. Overall these results support former work suggesting that an important proportion of the phages found in human gut are temperate phages with few lytic activity^10,12,46^.

To further study the question of active phages influences, we investigated MAGs growth rate using the software GRID that measure the dnaA (ori) / dif (ter) ratio^47^. We can clearly observe an inverse correlation between active phages and the growth rate of their hosts (Fig. 4a) suggesting that the more a phage is active, the less the bacterial population is growing. We then asked if some bacterial genera appeared more or less subject to phages attacks (Fig. 4b). Our results show that Bifidobacteriales and Selenomodales are less present in the group of targeted bacteria and, therefore, potentially less subject to phages lytic activity in the human gut microbiota.

### CrAss-like phages family in the human intestinal tract

CrAss-like phages are likely to become a large family of phages within the order Caudovirales and have been the subject of intensive research in the last years^13,48–50^. CrAss-like phages are wide spread in human gut and are predicted to infect bacteria from the Bacteroides phylum but, so far, only one fully identified host has been described “in vitro” from one crAss member, *Bacteroides intestinalis* 13. We searched for the presence of crAss-like phages in our different assemblies using a set of previously reported representatives phages of this family^49^ and detected 17 contigs, between 58 and 176 Kb, encoding proteins homologous to the crAss-like family proteins (Methods). Construction of a phylogenetic tree^45^ encompassing these 17 contigs as well as different representatives of the crAss-like family^49^ allowed us to find phages belonging to genera I, III, IV, VI and V (Fig. 5 and Supplementary Table 3). All these contigs were unambiguously attributed to MAGs belonging to the Bacteroides clade, as expected from previous studies^13,48,49^. We notably detect *Bacteroides vulgatus*, *Bacteroides uniformis, Bacteroides ovatus* and *Bacteroides thetaiotamicron* as hosts of these crAss-like phages, four species already suspected to be the host of crAss-like phages. Interestingly, we also detect unknown species from family Prevotellaceae and Porphyromonadaceae as host of other crAss-like phages. Some of these phages comes from the same sample and were attributed to the same MAG suggesting that crAss-like phages are highly abundant and potentially in competition in the human gut.

**Figure 5:**
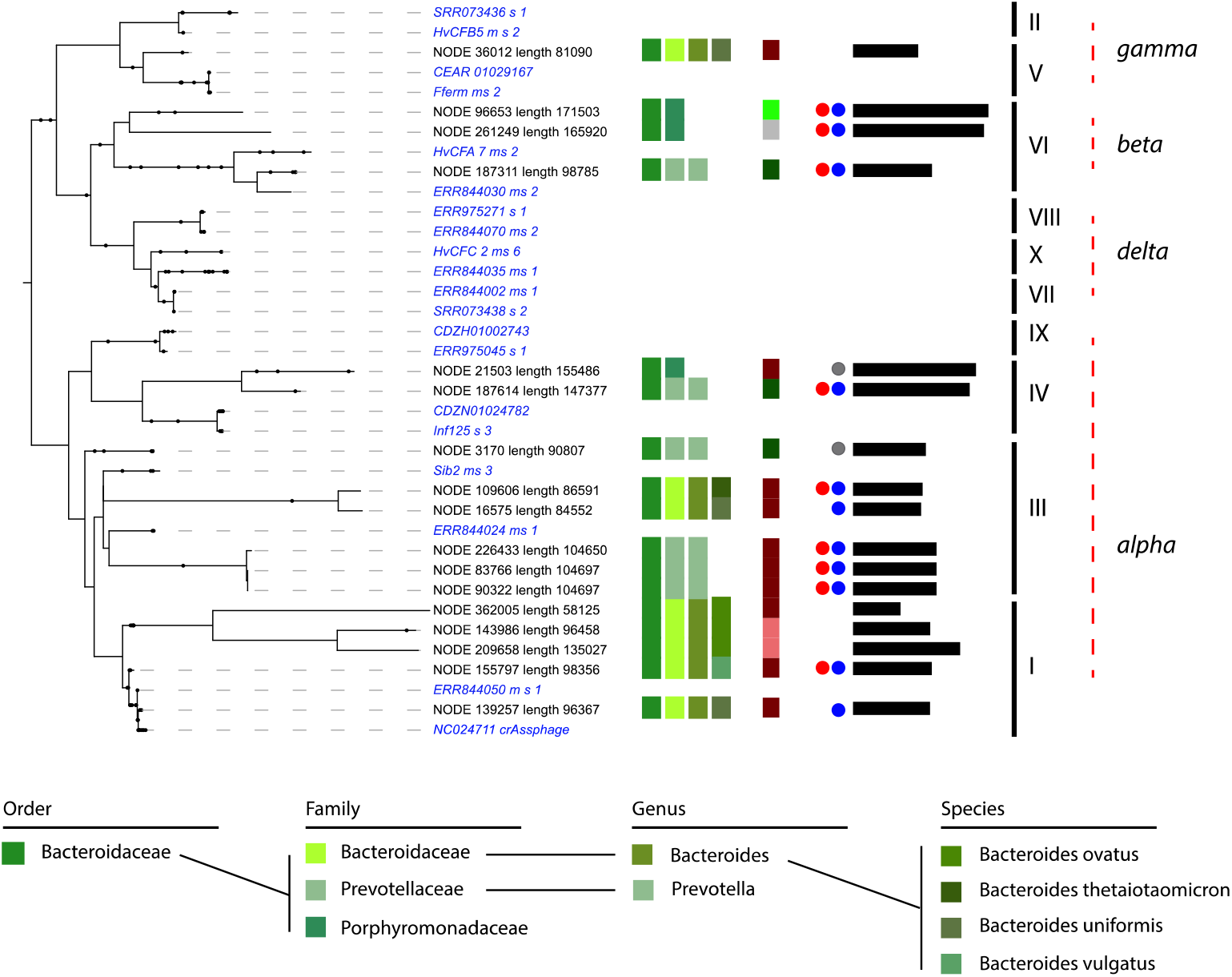
CrAss-like phages and their associated hosts. Phylogenetic tree of crAss-like phages contigs found in the 10 assemblies. Representatives of the 10 genera of crass-like phages described by Guerin *et al*. are included (in *italic blue*). Names of the different contigs are indicated on the left of each branch. Vertical colored stripes indicate, from left to right: host MAG order, host MAG family, host MAG genus, host MAG species, ratio of the read coverage of phages vs. read coverage of MAG (red – ratio > 2; light red – 2 > ratio > 1.25; grey – 1.25 > ratio > 0.75; light green – 0.75 > ratio > 0.5; dark green – ratio < 0.5). Red dot: circular signal characterized by VirSorter. Blue and grey dot: strong and weak circular signal characterized by 3C, respectively. Black lines: size of the contigs (min = 58,125 pb; max = 171,503 bp). Genera of the crass-like phages are indicated on the right of the tree.

Among this 17 contigs, 9 are predicted to be circular by VirSorter. We confirmed this topology using the 3D signal (Fig. 5 and Supplementary Fig. 9). We identified one contig as circular based on this 3D contact and detected a weak circular signal for two others, demonstrating that 3C methods can be used for topological characterization. Finally, we also computed the phage – host ratio for these 17 contigs and demonstrate that crAss-like phages are a very active family of phages (12 contigs exhibit a ratio ≥ 1.25). On the other hand, we could not confirm that these ratios are linked to a low growth rate and further analysis will be needed to understand the impact of crAss-like phages on their hosts. All in all, our results improved significantly our understanding of the spreading of the ubiquitous but elusive crAss-like family, while confirming its link with Bacteroides.

## Discussion

Bacteriophages, the viruses that infect bacteria, are the most abundant biological entities on earth and have a major impact on microbial communities^51^. Development of high-throughput DNA sequencing technologies and its application on environmental samples have allowed to bypass virus isolation and have revealed the existence of a wide variety of phages in all ecosystems^48,52,53^. With the increasing interest in studying human gut microbiota, several studies and computational developments have shed light on the composition and dynamics of its viral part^11,54,55^. These approaches have notably allowed the description of completely new phages family such as the ones from the crAss-like family^48–50^. However, methods allowing to study phages and bacteria concomitantly are still in their infancies^12,19^. Moreover, most of the metagenomic approaches are based on the use of multiple samples and there is an increasing need to be able to study single samples. Proximity ligation assay applied to metagenomic samples (meta3C, metaHiC) are promising approaches to study *in situ* microbial communities as a whole starting from single samples^19,20^. In the present study, we applied those approaches to ten human gut samples, characterizing hundreds of high-quality bacterial genomes (MAGs) as well as phage sequences, and revealing the large multiscale network of interactions between them.

Our study confirms that most of the phages are quite specific to their host and do not shift easily to another host, at least in the human intestinal tract^43,44^. On the other hand, nearly 1/3rd of the identified phage contigs exhibit contacts with more than one MAGs. Analysis aiming at characterizing those relationships are needed to better understand their role in the regulation of the balance of the human gut.

The phage-host ratio computed in this study tends to show that phages appear to be slightly more abundant than bacteria in human intestinal tract but nothing comparable to other environments^56^. Therefore, our work confirms findings that majority of the phages present within human gut are temperate phages with no or few lytic activity^46^. This result is in adequation with previous studies that showed a limited phage-host dynamic and a relative homeostasis of this environment with few examples of a phage overrepresented compared to its hosts as for the crAss phage. On the other hand, we also found nearly 10% of the phage population exhibiting a ratio over 2 compared to their hosts and representing actively replicating phages. This population could play an important role in the maintenance of bacterial population and will need further analysis. Notably, we found that members from the Bifidobacteriales and the Selenomonadales clades appear to be less subject to phages lytic activity.

As expected from their overrepresentation in the human gut population^57^, we found several candidate members of the crAss-like phages family in our samples. This have allowed us to characterize the host of 17 crass-like contigs, all belonging to different members of the Bacteroides clade as previously predicted^48^. This represents a major breakthrough in the field as many people have tried to cultivate those phages and their hosts *in vitro* since their discovery in 2014 with only one success^13^. This finding highlights the potential of our approach which is not expensive and is easily applicable to human gut sample. We conclude that crAss-like phages constitute a large family of phages able to infect various members of the Bacteroides clade. Application of meta3C on larger cohorts will allow, in the future, to better understand relation between crAss phages and this clade as well as the specificity of each phages sub-group to their hosts. CrAss-like phages family harbor different features of phage and plasmid and generally encode proteins allowing their replication. The concept of phasmid or phage plasmid 58 is not new but we hypothesize that crAss-like family belong to this type of genetic elements and are widely distributed in the Bacteroides clade.

No doubt that future studies, coupling meta3C with different approaches on larger human cohort, will shed light and new finding about relationships between phages, plasmids and bacteria. For instance, by coupling meta3C approaches with Virus Like Particles (VLP) purification, future studies will offer an unprecedent view of phages-bacteria relationships and will bring us to a better understanding of this particular and important ecosystem.

## Agreement

This research receives the agreement n°N18 from Institut Pasteur (ICAReB).

## Acknowledgment and funding

This research was supported by funding to R.K. from the European Research Council under the Horizon 2020 Program (ERC grant agreement 260822) and JPI-EC-AMR STARCS ANR-16-JPEC-0003-05.

We thank all our colleagues from the laboratory, especially Pierrick Moreau and Lyam Baudry for discussions, feedback, and comments. We also Thank M.A. Petit and A. Laffitte for constructive comments on the data analysis and the manuscript.

## Authors contributions

Experiments: AT and MM; Data analysis: MM; Conceptualization: MM and RK; Writing: MM and RK; Funding acquisition: RK

## Table of contents

**Supplementary Fig. 1:** Comparisons of 3C, Hi-C and eHi-C protocols.

**Supplementary Fig. 2:** Comparisons of MAGs obtained for the sample #9010 using 3C or eHi-C.

**Supplementary Fig. 3:** Phylogenetic tree for the 10 processed samples.

**Supplementary Fig. 4:** Comparisons of MAGs taxonomic abundancy and reads taxonomic abundancy.

**Supplementary Fig. 5:** Relation between abundancy and completion / contamination for the retrieved MAGs.

**Supplementary Fig. 6:** Classification of phages contigs.

**Supplementary Fig. 7:** Comparison of phages and their characterized hosts between samples.

**Supplementary Fig. 8:** Phages – hosts ratio.

**Supplementary Fig. 9:** CrAss-like phages contact maps.

**Supplementary table 1:** Assembly statistics.

**Supplementary table 2:** Mapping statistics and 3D ratio.

**Supplementary table 3:** CrAss-like phages contigs.

**Supplementary Figure 1.**
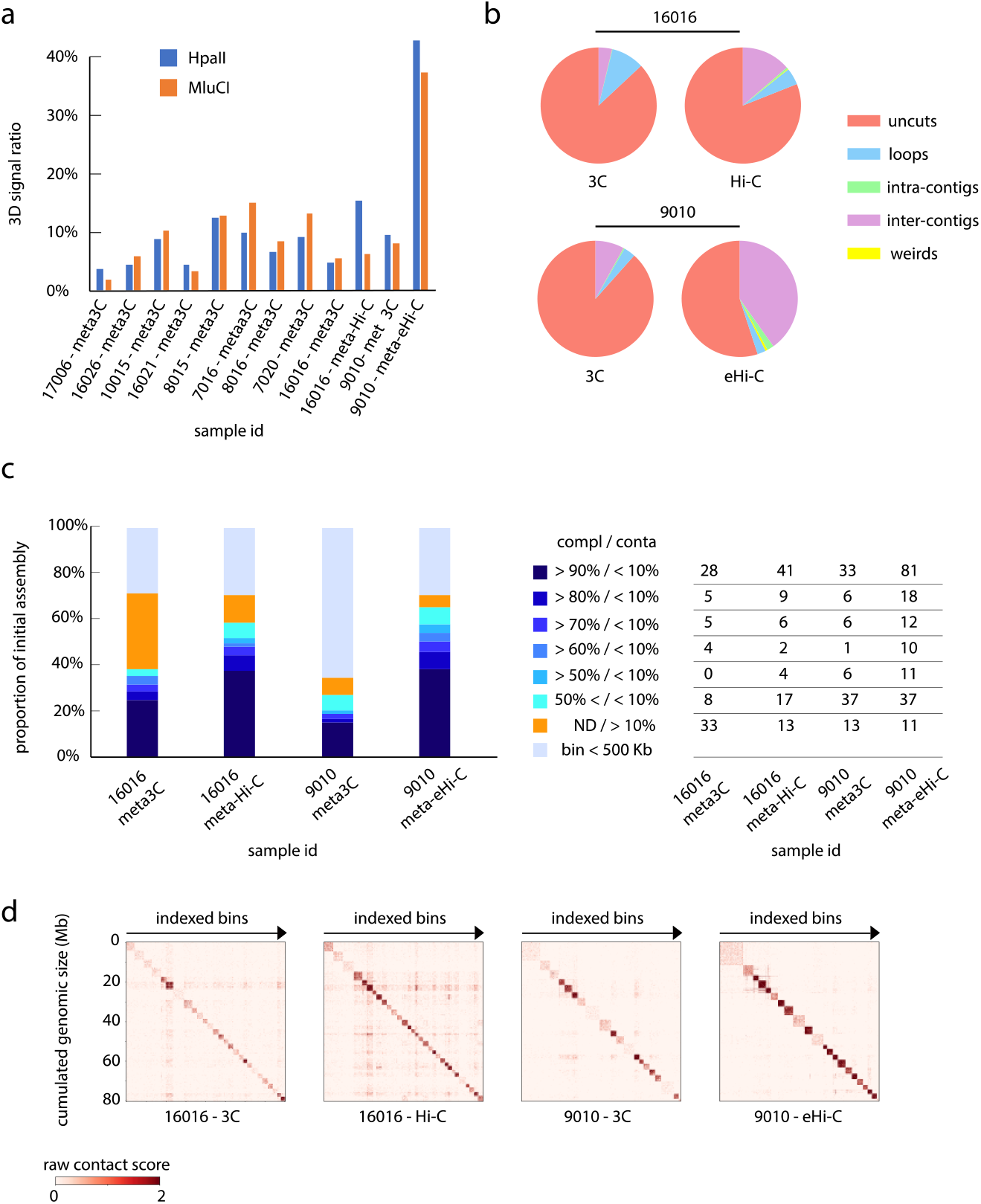
Comparisons of 3C, Hi-C and eHi-C protocols. **a.** Bar plot of 3D signal ratio calculated for each constructed library. The 3D ratio is defined as the ratio of the PE reads that link unambiguously two contigs together compared to the number of mapped PE reads (see Methods). **b.** Pie charts indicating the proportion of the different PE reads types for the libraries 16016_3C, 16016_Hi-C, 9010_3C and 9010_eHi-C1. Red: uncuts, Blue: loops, Green: intra-contigs interaction, Purple: inter-contigs interaction, Yellow: weird [weird= a PE reads that interacts with the same restriction fragment]. **c.** To compare the data recovered with the different protocols, we initialized MetaTOR with equal number of PE-reads from the libraries generated from the sample #16016 (50 million PE reads; 3C and Hi-C libraries) and #9010 (100 million PE reads; 3C and eHiC libraries) and their corresponding assemblies. (left) bar charts show the proportion of the different bins obtained using 3C, Hi-C or eHi-C on the samples 16016 and 9010 and starting from the same numbers of PE reads. Colors indicates the quality of the bins as indicated on the middle. (left) number of MAG in the different categories. The mean completion and contamination rates were 41.1 % and 4.3 % for 16016-3C; 56.9 % and 6.7 % for 16016-Hi-C; 68.3 % and 5.4 % for 9010-3C; 75.9 % and 4.2 % for 9010-eHi-C. The number of bins exhibiting less than 10% of contamination is strongly increased for the eHi-C method. **d.** Contact maps of the largest bins recovered after 100 Louvain iterations and 10 recursive Louvain iterations (1 pixel = 40 kb). The x and y axes are labeled with the cumulated DNA size and the index of the community, respectively.

**Supplementary Figure 2.**
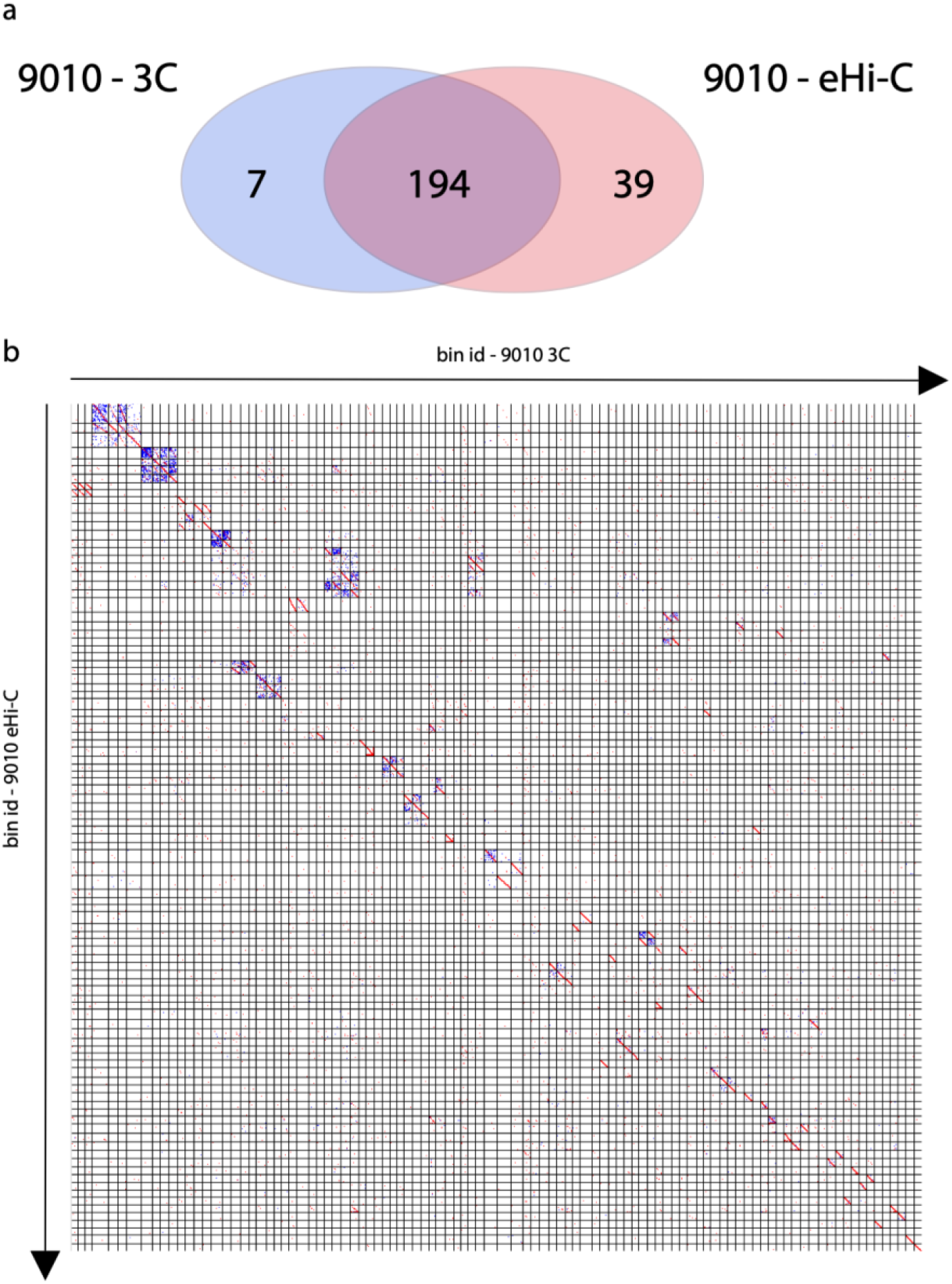
Comparisons of MAGs obtained for the sample #9010 using 3C or eHi-C. **a.** Venn diagram of the MAGs obtained using 3C or eHi-C data. We took all the reads from the #9010 3C libraries (HpaII and MluCI) on one hand and all the reads for the #9010 eHi-C libraries on the other hand and aligned them on the reference assembly to generate the two networks. These networks where processed with MetaTOR. We then compared the 233 bins > 500 kb retrieved for the eHi-C protocol with the 201 bins > 500kb recovered from the meta3C protocol. 194 bins (96% of the lowest number) exhibit more than 80 % of identity, showing that both approaches are highly concordant despite the slightly noisier signal of the meta3C method, which can be overcome by a higher sequencing depth. **b.** LAST alignment of the MAGs > 1Mb retrieved using 3C or eHi-C on sample #9010. X and Y-axis indicate the MAGs index for 3C (X) or eHi-C (Y).

**Supplementary Figure 3.**
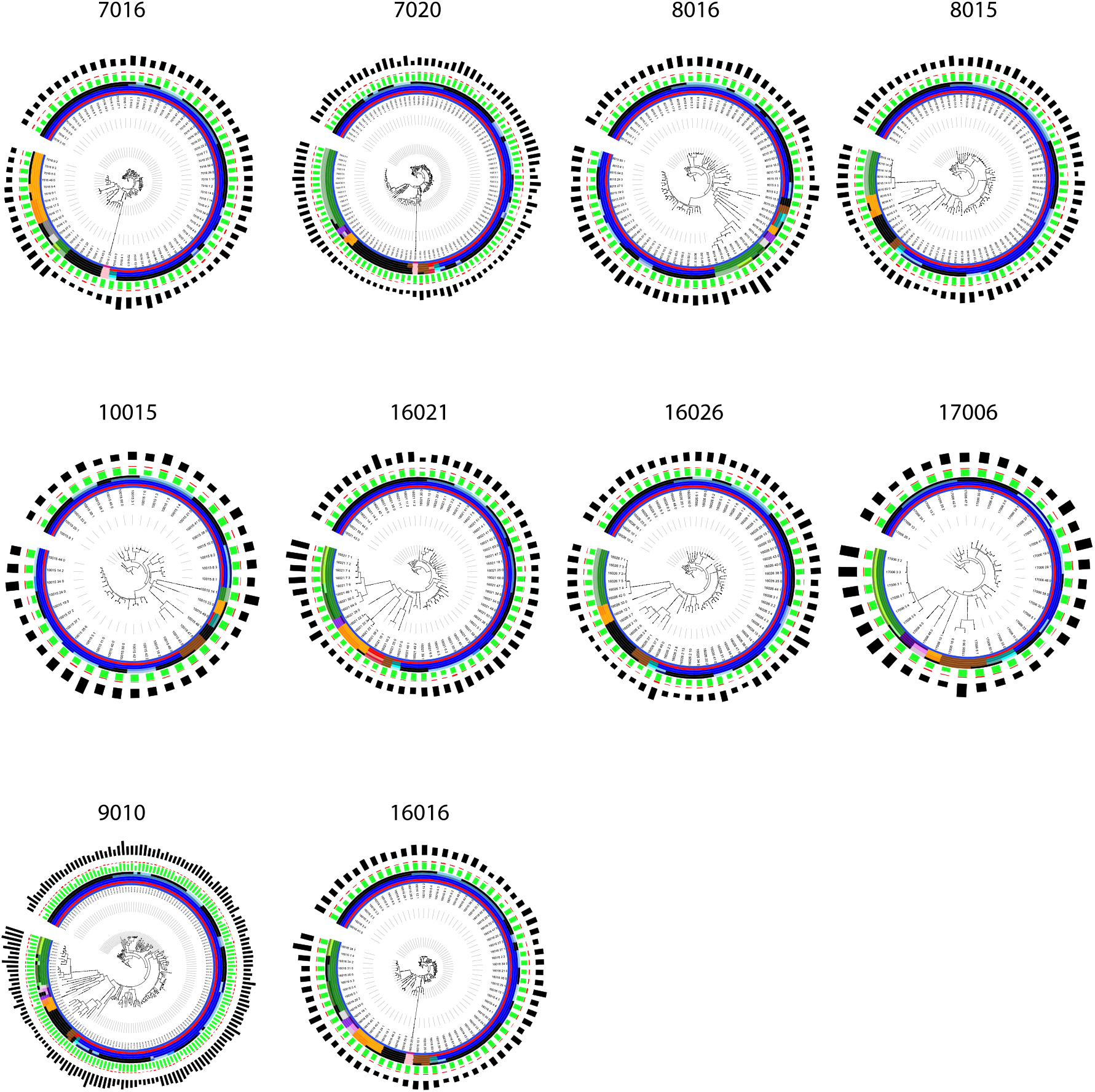
phylogenetic tree for the 10 processed samples. Only MAGs with a completion above 50% and a contamination below 20% are integrated in each tree. Colors of the inner six layers indicate the taxonomy of the MAG attributed by CheckM (from center to periphery: Phylum, Class, Order, Family, Genus, species). Green and red bars in the following layers indicated completion and contamination. Black bars in the outer layer indicated MAG size.

**Supplementary Figure 4.**
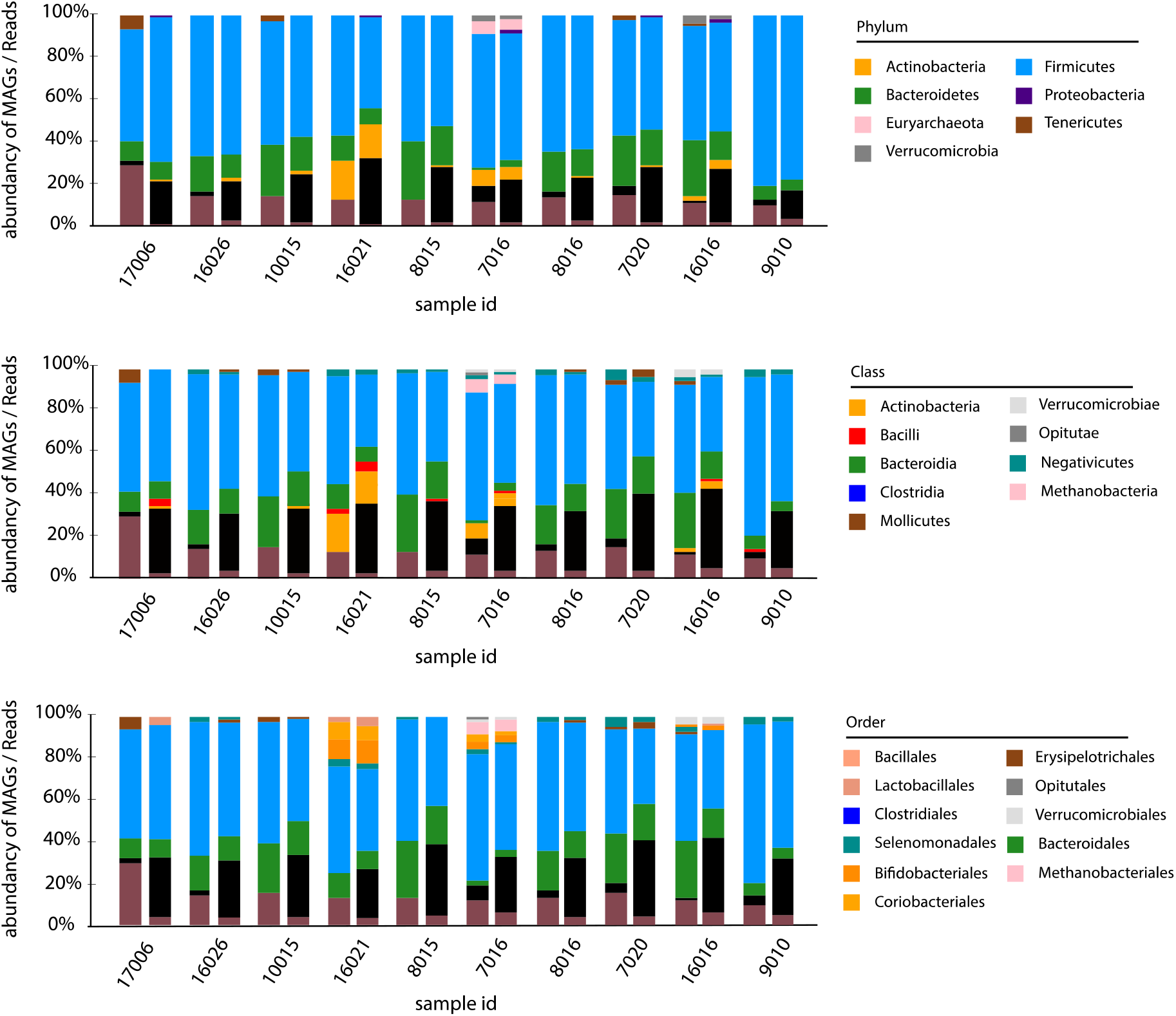
comparisons of MAGs taxonomic abundancy and reads taxonomic abundancy. Bar charts of MAGs taxonomic abundancy (left) and reads taxonomic abundancy using Kaiju2 (right) for each sample. Different taxonomic levels are shown (upper: Phylum; middle: Class; bottom: Order).

**Supplementary Figure 5.**
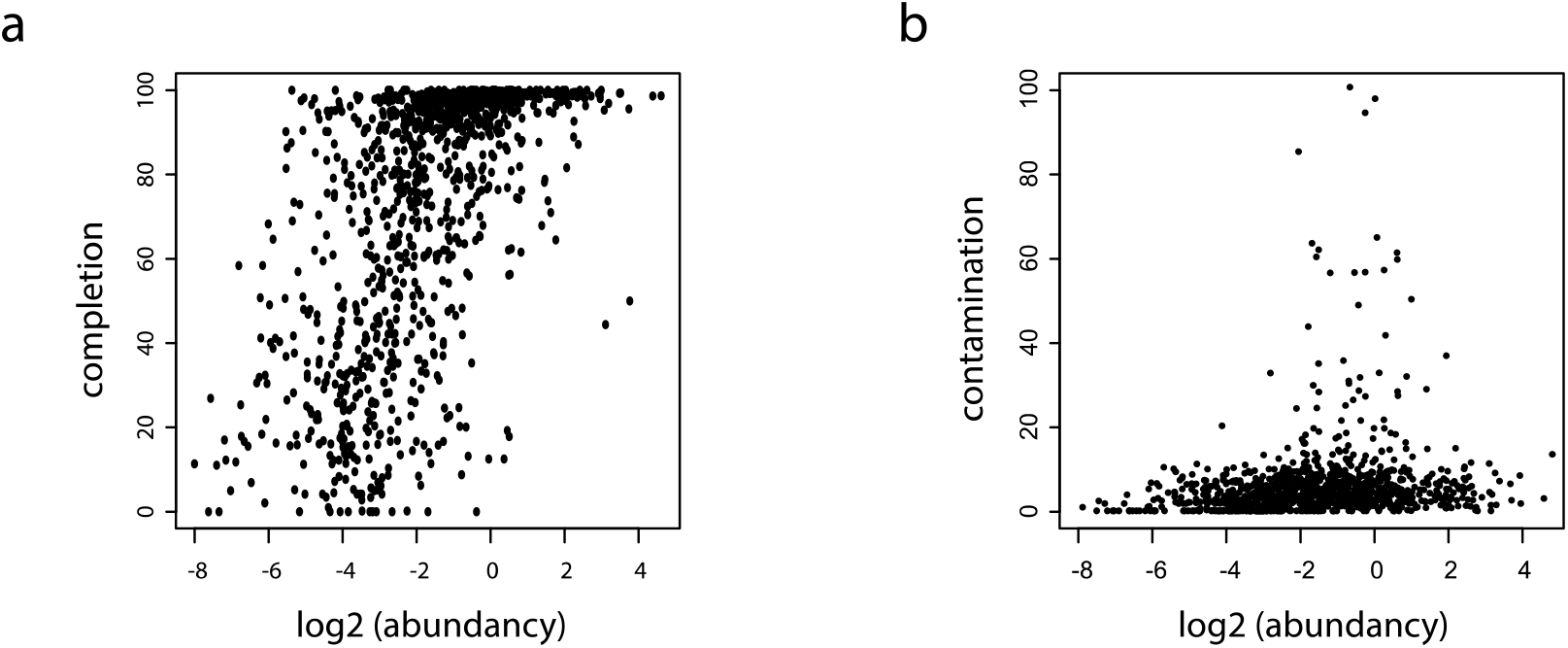
Relation between abundancy and completion / contamination for the retrieved MAGs. **a.** Plot of the completion (Y-axis) in function of abundancy(log2) (X-axis) for the retrieved MAGs (i.e. bins > 500 Kb). **b.** Plot of the contamination (Y-axis) in function of abundancy(log2) (X-axis) for the retrieved MAGs (i.e. bins > 500 Kb).

**Supplementary Figure 6.**
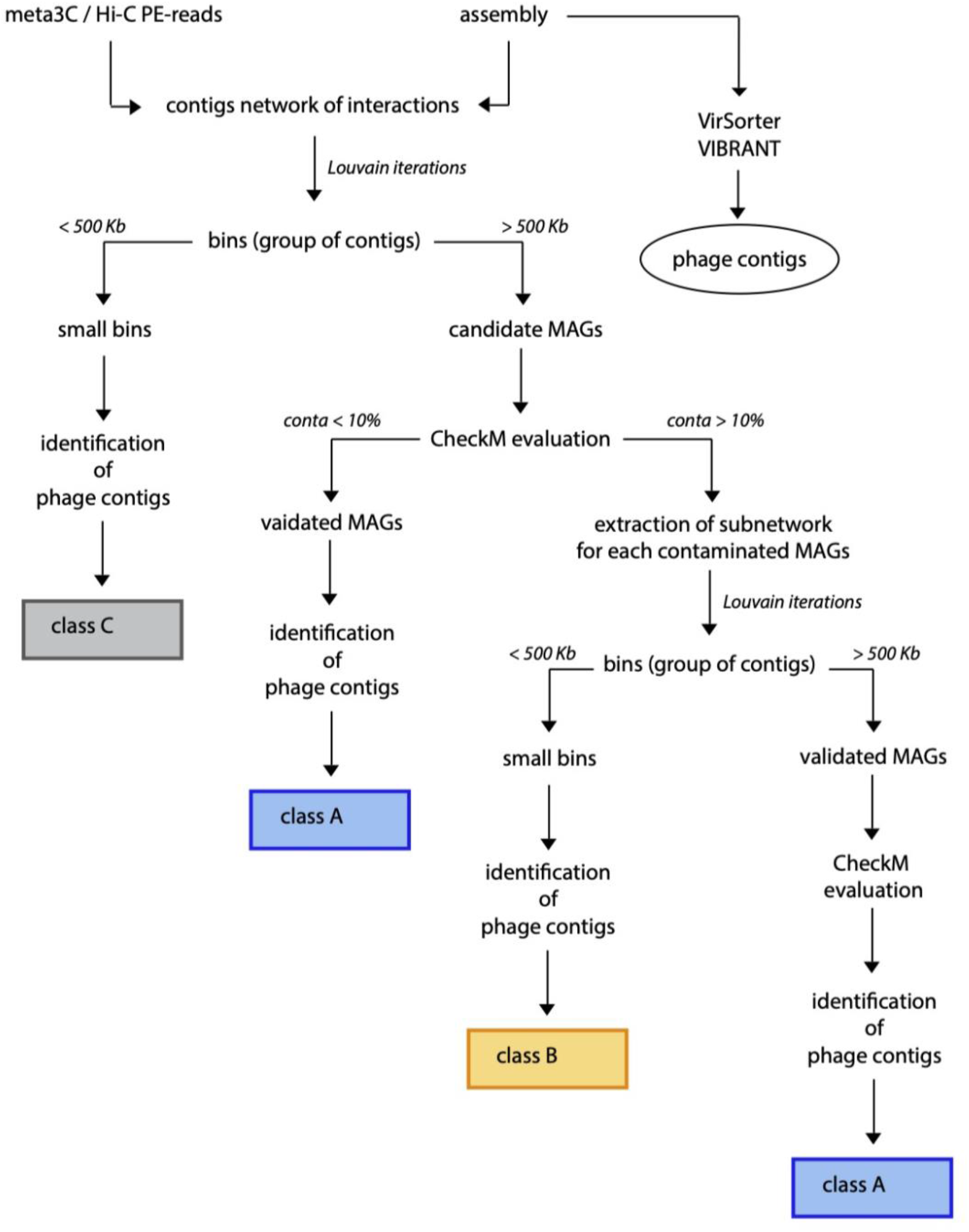
Classification of phages contigs. Scheme of the followed process to classify phage contigs in the 3 classes (A,B,C) (see Methods).

**Supplementary Figure 7.**
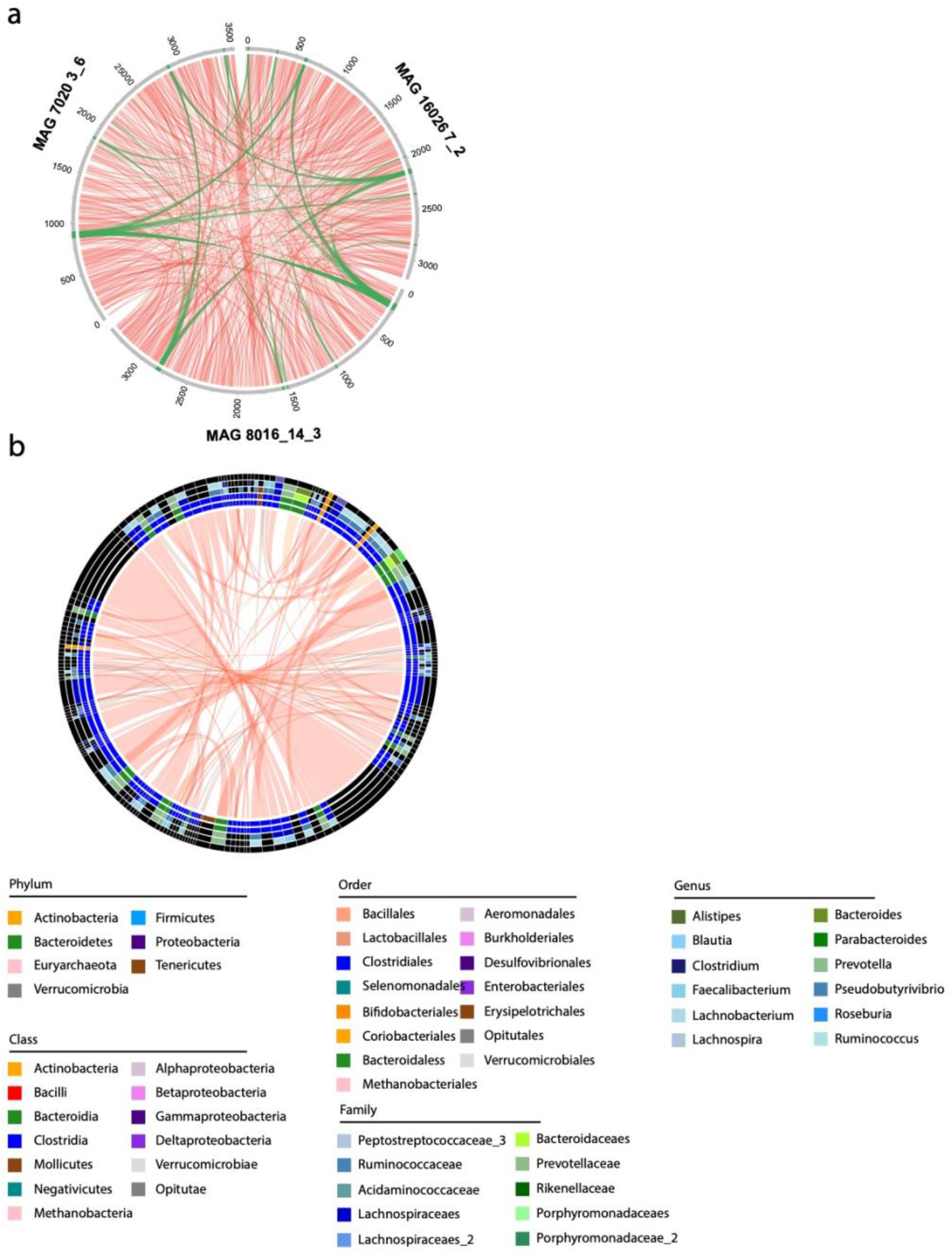
Comparison of phages and their characterized hosts between samples. **a.** Circos representation of 3 homologous MAGs from samples #7020, #16026 and #8016. MAGs are indicated in grey with their respective size indicated in Kilobases. Phages contigs are indicated in green. Red Links represents blast hits with an identity above 90%. Green links represents blast hits with an identity above 90% for identified phages contigs. **b.** Circos representation of the different phages belonging to the same Genus characterized in the 10 samples. Phages are defined as belonging to the same Genus if their shared at least 70% identity over 70% of their length. The different circles represent the different characterized hosts at different taxonomic levels (from inside to outside: Phylum, Class, Order, Family, Genus). Links are colored in function of agreement between hosts taxnomy: Red – same taxonomic annotations at the genus level; Orange – same taxonomic annotations at the Class level; Grey – different annotations at the Phylum level. Legend for taxonomic annotations is indicated under the circos.

**Supplementary Figure 8.**
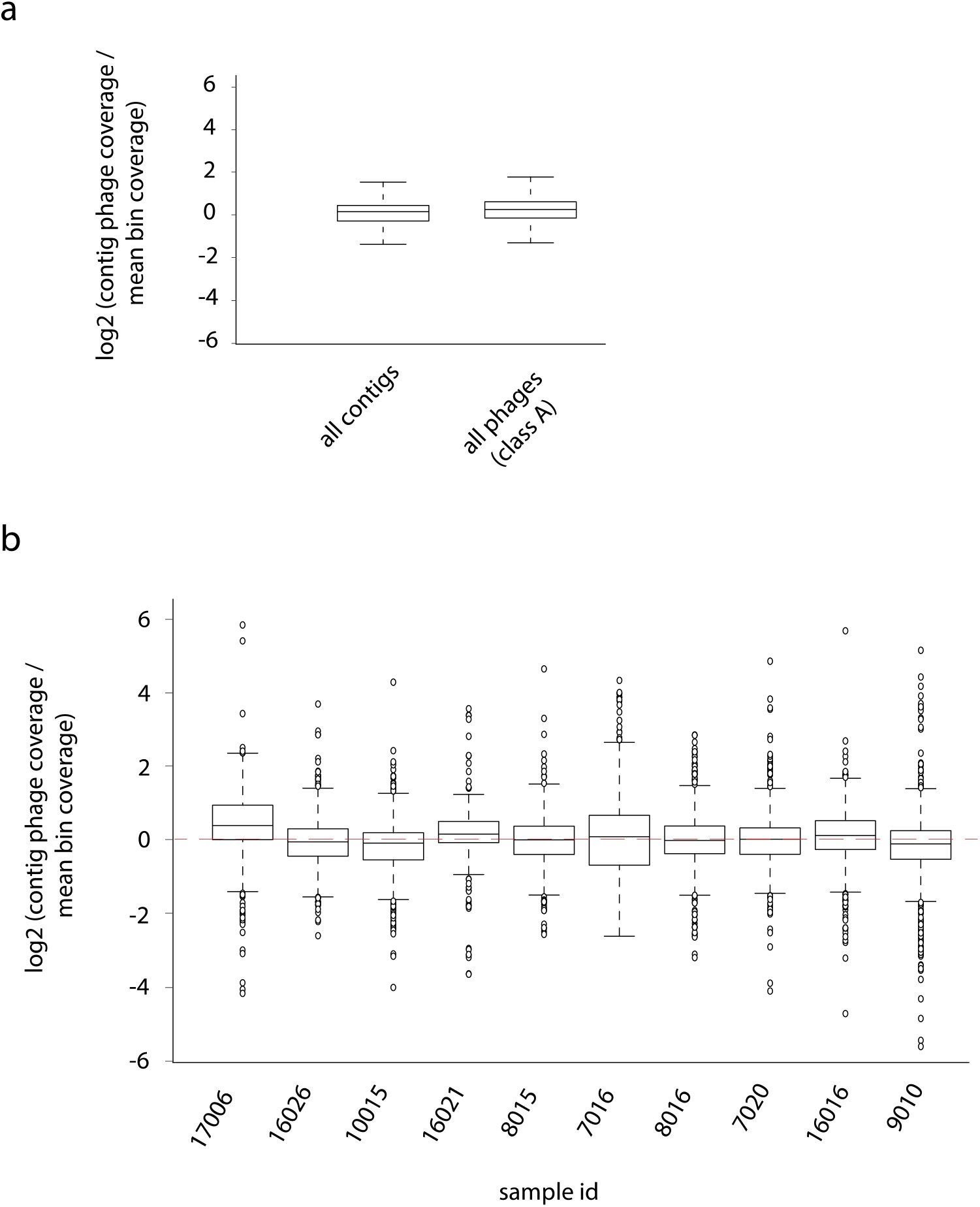
phages – hosts ratio. **a**. boxplot of the coverage ratio between contigs and their associated MAG (log2 scale) for all contigs unambiguously associated to a MAG (left) and phage contigs from class A (right). **b**. Boxplot of the coverage ratio between phage contigs and their associated MAG (log2 scale) for the 10 samples. Dashed red line indicate a ratio of 1.

**Supplementary Figure 9.**
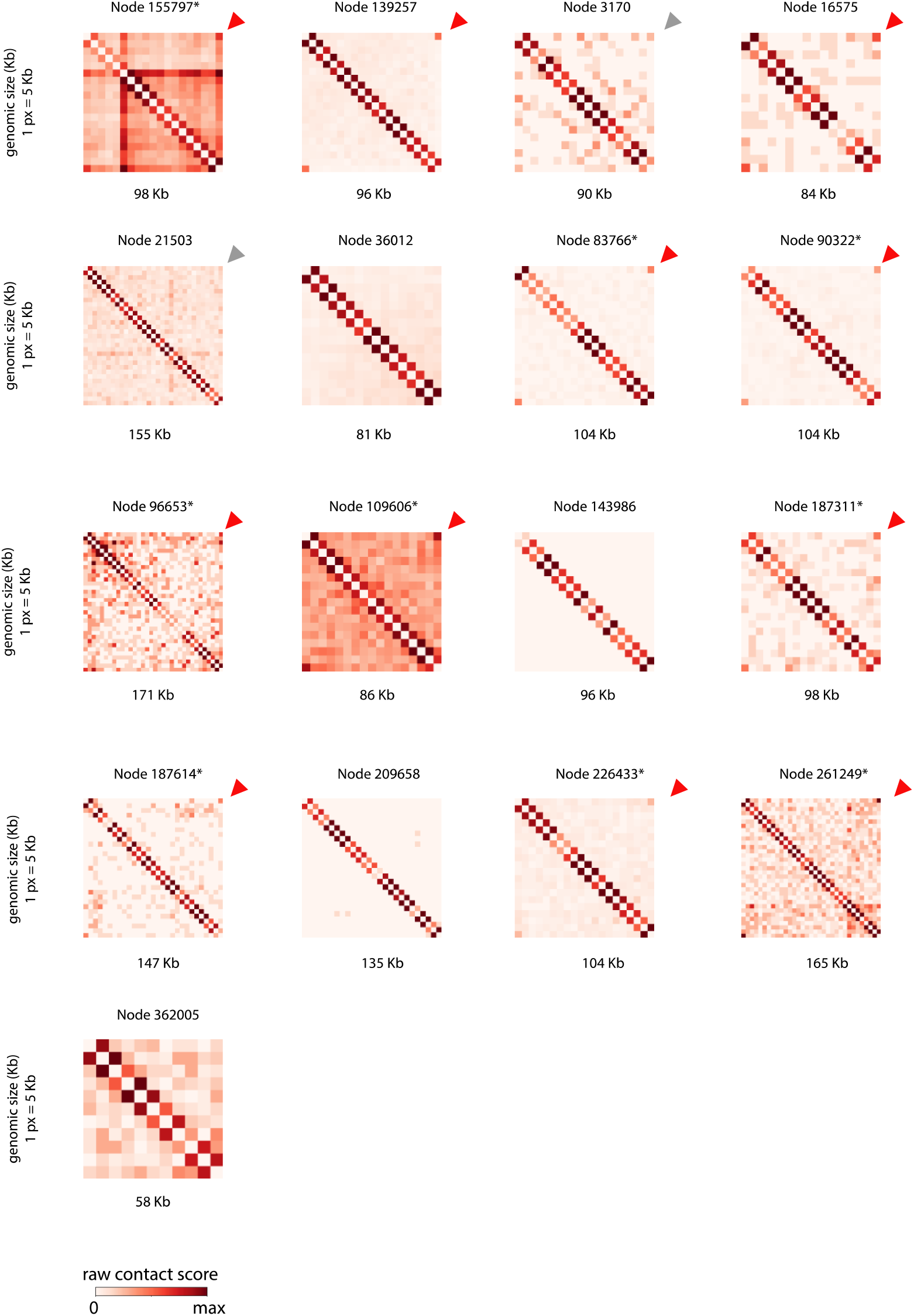
CrAss-like phages contact maps. Raw contact matrices of the different crAss-like phage contigs found in the 10 assemblies. Each contact map is displayed using 5Kb bins (1 pîxel = 5 Kb). Arrow in the upper right corner indicates a 3C circular signal (red = strong signal, grey = weak signal, no arrow = no signal). Contig ID and size are indicated upper and under each matrix. Star in the contig ID indicates that VirSorter characterize this contigs as circular.

**Supplementary Table 1.** Assembly statistics.

| Id | sex | age | enzyme | raw PE-reads<br>(2x75 bp) | filtered PE-reads | assembly (contigs > 500 bp) |  |  |
| --- | --- | --- | --- | --- | --- | --- | --- | --- |
|  |  |  |  |  |  | size (bp) | contigs | N50 (bp) |
| 17006 | F | 19 | HpaII | 69 115 370 | 64 215 120 | 242 050 174 | 96 872 | 8 631 |
|  |  |  | MluCI | 58 557 235 | 55 067 270 |  |  |  |
| 16026 | M | 20 | HpaII | 62 344 476 | 57 625 084 | 299 556 023 | 132 589 | 4 642 |
|  |  |  | MluCI | 66 017 028 | 61 742 909 |  |  |  |
| 10015 | F | 26 | HpaII | 48 226 880 | 46 076 874 | 228 816 554 | 99 416 | 4 674 |
|  |  |  | MluCI | 45 253 636 | 43 440 164 |  |  |  |
| 16021 | F | 28 | HpaII | 88 144 160 | 83 223 643 | 321 521 194 | 116 351 | 8 066 |
|  |  |  | MluCI | 105 816 172 | 96 257 997 |  |  |  |
| 8015 | F | 64 | HpaII | 47 079 274 | 45 855 674 | 318 332 624 | 137 809 | 5 031 |
|  |  |  | MluCI | 46 985 066 | 45 988 957 |  |  |  |
| 7016 | M | 64 | HpaII | 48 666 658 | 47 158 690 | 339 356 582 | 166 960 | 3 857 |
|  |  |  | MluCI | 49 606 315 | 48 419 453 |  |  |  |
| 8016 | F | 69 | HpaII | 71 046 963 | 69 086 479 | 337 623 345 | 143 613 | 5 535 |
|  |  |  | MluCI | 61 331 543 | 59 687 922 |  |  |  |
| 7020 | M | 73 | HpaII | 53 137 532 | 51 450 802 | 430 993 887 | 199 668 | 4 126 |
|  |  |  | MluCI | 74 379 917 | 72 428 112 |  |  |  |
| 16016 | F | 35 | HpaII | 59 333 139 | 55 876 445 | 314 197 889 | 140 409 | 5 418 |
|  |  |  | MluCI | 72 673 730 | 68 783 443 |  |  |  |
| 9010 | F | 60 | HpaII | 210 604 009 | 206 957 360 | 617 881 257 | 251 469 | 5 600 |
|  |  |  | MluCI | 202 789 023 | 198 771 314 |  |  |  |

**Supplementary Table 2.** Mapping statistics and 3D ratio.

|  |  | ratio (mapped PE-reads on different contigs / raw PE-reads) |  |  |  |  |  |  |  |
| --- | --- | --- | --- | --- | --- | --- | --- | --- | --- |
|  |  | HpaII (or Sau3AI) |  |  |  | MluCI |  |  |  |
| id | protocol | initial PE-reads | mapped PE-reads (no ambiguous, MQ >= 20) | inter-contig contacts | ratio | raw PE-reads | mapped PE-reads (no ambiguous, MQ >= 20) | inter-contig contacts | ratio |
| 17006 | 3C | 64215120 | 52188432 | 1814822 | 3,48% | 55067270 | 45819103 | 877448 | 1,92% |
| 16026 | 3C | 57625084 | 44338605 | 1865567 | 4,21% | 61742909 | 46525292 | 2683307 | 5,77% |
| 10015 | 3C | 46076874 | 36518505 | 3068991 | 8,40% | 43440164 | 34626773 | 3407767 | 9,84% |
| 16021 | 3C | 83223643 | 68588525 | 2965637 | 4,32% | 96257997 | 79930032 | 2670294 | 3,34% |
| 8015 | 3C | 45855674 | 35854912 | 4299603 | 11,99% | 45988957 | 35853789 | 4403148 | 12,28% |
| 7016 | 3C | 47158690 | 33967366 | 3246134 | 9,56% | 48419453 | 34058489 | 4966086 | 14,58% |
| 8016 | 3C | 69086479 | 53997499 | 3449134 | 6,39% | 59687922 | 45886287 | 3692661 | 8,05% |
| 7020 | 3C | 51450802 | 39341412 | 3416173 | 8,68% | 72428112 | 53064475 | 6717028 | 12,66% |
| 16016 | 3C | 55876445 | 41174584 | 1845660 | 4,48% | 68783443 | 51429206 | 2648451 | 5,15% |
|  | Hi-C | 50594809 | 36616746 | 5461001 | 14,91% | 83950053 | 62699625 | 3729559 | 5,95% |
| 9010 | 3C | 206957360 | 169153016 | 15305524 | 9,05% | 198771314 | 161725492 | 12284430 | 7,60% |
|  | eHi-C | 102203764 | 89489651 | 36891289 | 41,22% | 113637817 | 78267233 | 28188719 | 36,02% |

**Supplementary Table 3.**
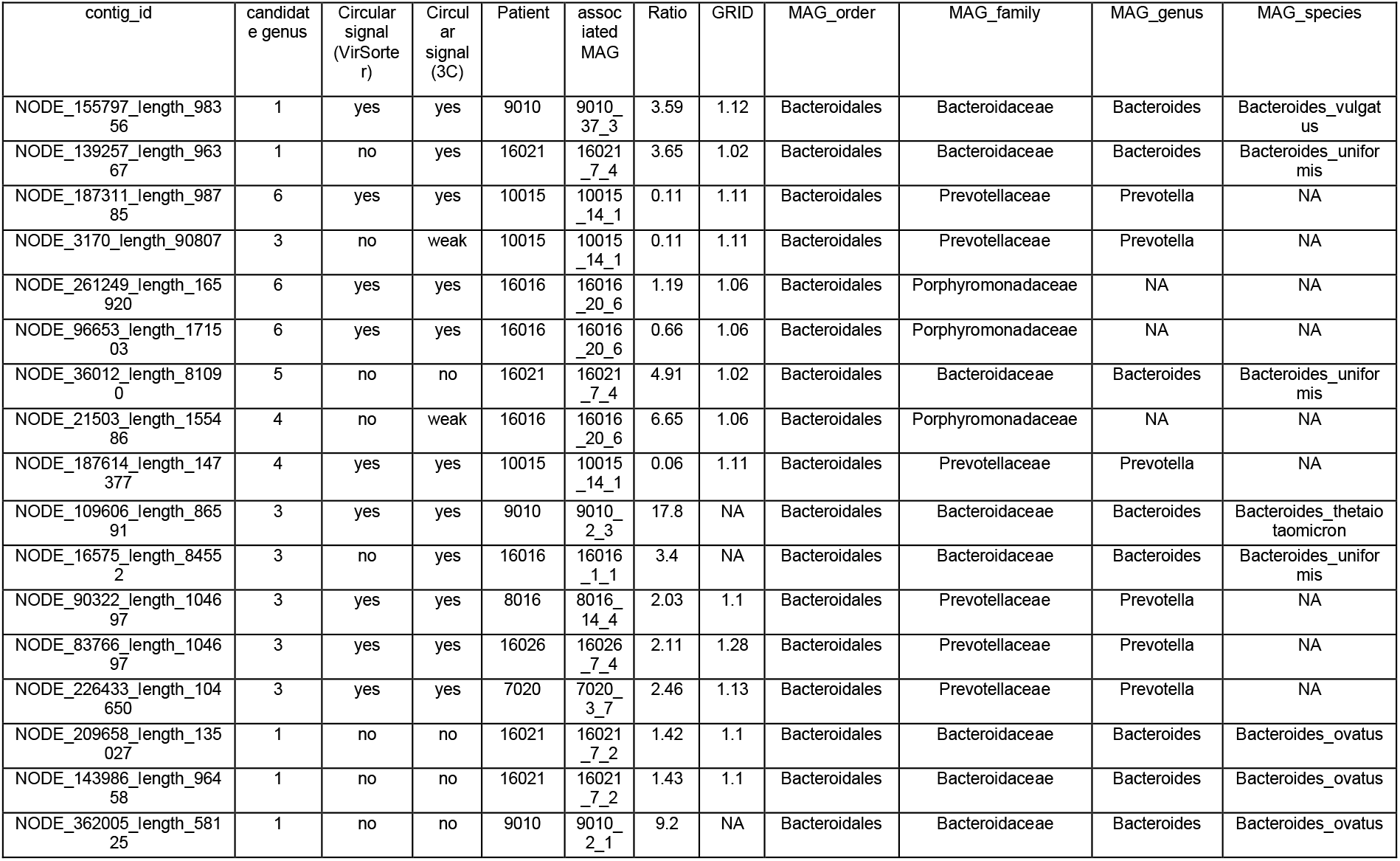
CrAss-like phages contigs.

